# Phase Unwrapping with a Rapid Opensource Minimum Spanning TreE AlgOrithm (ROMEO)

**DOI:** 10.1101/2020.07.24.214551

**Authors:** Barbara Dymerska, Korbinian Eckstein, Beata Bachrata, Bernard Siow, Siegfried Trattnig, Karin Shmueli, Simon Daniel Robinson

## Abstract

**Purpose:** To develop a rapid and accurate MRI phase unwrapping technique for challenging phase topographies encountered at high magnetic fields, around metal implants or post-operative cavities, that is sufficiently fast to be applied to large group studies including Quantitative Susceptibility Mapping and functional MRI (with phase-based distortion correction).

**Methods:** The proposed path-following phase unwrapping algorithm, ROMEO, estimates the coherence of the signal both in space - using MRI magnitude and phase information - and over time, assuming approximately linear temporal phase evolution. This information is combined to form a quality map that guides the unwrapping along a three-dimensional path through the object using a computationally efficient minimum spanning tree algorithm. ROMEO was tested against the two most commonly used exact phase unwrapping methods: PRELUDE and BEST PATH in simulated topographies and at several field strengths: in 3 T and 7 T in vivo human head images and 9.4 T ex vivo rat head images.

**Results:** ROMEO was more reliable than PRELUDE and BEST PATH, yielding unwrapping results with excellent temporal stability for multi-echo or multi-time-point data. ROMEO does not require image masking and delivers results within seconds even in large, highly wrapped multi-echo datasets (e.g. 9 seconds for a 7 T head dataset with 31 echoes and a 208 x 208 x 96 matrix size).

**Conclusion:** Overall, ROMEO was both faster and more accurate than PRELUDE and BEST PATH delivering exact results within seconds, which is well below typical image acquisition times, enabling potential on-console application.

## INTRODUCTION

The complex signal in Magnetic Resonance Imaging (MRI) can be divided into two constituents: magnitude (*M*) and phase (*θ*). MRI phase is proportional to local deviations in the static magnetic field, Δ*B*_0_ (Hz), via the relation *θ*~2*πTE* · Δ*B*_0_, where TE is the echo time. Knowledge of Δ*B*_0_ can be used to correct image distortions (1,2), visualize veins and microbleeds using Susceptibility Weighted Imaging (SWI) (3), assess iron-rich tissues or calcifications via Quantitative Susceptibility Mapping (QSM) (4), and to estimate blood flow (5) or temperature changes in tissue (6). The measured phase, φ, is a projection of the true phase *θ* into the 2π range. This gives rise to abrupt changes, i.e. wraps, which do not represent the spatial and temporal continuity of *θ* within the object and require unwrapping.

A questionnaire completed by 46 participants at the 5^th^ International Workshop on MRI Phase Contrast and Quantitative Susceptibility Mapping in South Korea (September 2019) indicated that 84.8 % of participants use Laplacian unwrapping (7) in their work, 32.6% use PRELUDE (8), 30.4 % use BEST PATH (9) and 2.2% use Graph-Cut (10) (unpublished results reported by Prof. Peter C.M. van Zijl of Johns Hopkins University). Laplacian unwrapping is the most robust method currently available, providing globally smooth phase results (i.e. no abrupt jumps) within seconds even for large datasets with low signal-to-noise ratio (SNR), explaining its popularity in QSM. However, it does not yield exact results for *θ*, which makes it unsuitable for applications such as distortion correction, flow or temperature measurements. Moreover, Laplacian unwrapping introduces large phase variations around regions with sharp phase changes, such as veins, which corrupts QSM results around these structures (11–14). PRELUDE and BEST PATH are the methods of choice when exact phase results are desired. They assume that phase changes between voxels which exceed π are indicative of wraps. PRELUDE belongs to the class of region-growing spatial unwrapping approaches, which divide the volume into wrapless regions, i.e. groups of contiguous voxels containing ranges of values which are less than π, and assess phase changes at the borders between them. PRELUDE is a relatively robust algorithm considered to be the gold standard (11) but can take several hours or even days to unwrap large datasets with challenging phase topographies. A substantial reduction in computation time has been achieved using a recently developed method based on PRELUDE, called SEGUE (15), by simultaneously unwrapping and merging multiple regions. However, SEGUE can still take more than 10 minutes to unwrap more challenging datasets (e.g. 17 m 35 s ± 9m 26 s using a 3.5 GHz processor for images acquired at 3T with matrix size = 220 x 220 x 240 and TE = 18.9 ms) making potential on-console implementation impractical. Path-following approaches, such as BEST PATH (9), usually provide solutions within seconds even for highly wrapped images with large matrix sizes (11). This can be particularly useful in large studies including functional MRI, where hundreds of 3D image volumes are often acquired per subject. Path-following algorithms compare the phase in adjacent voxels, beginning at one location and proceeding to neighbouring voxels in an order dictated by the reliability of the information in the voxels and how well they are connected, i.e. a quality map. BEST PATH is rapid but more prone to errors than PRELUDE, especially in regions where a corresponding magnitude image has low SNR. For a comprehensive comparison of phase unwrapping algorithms, we refer the reader to Refs (11) and (16).

To overcome the shortcomings of the exact phase unwrapping algorithms currently available, we propose a new path-following algorithm called ROMEO: <u>R</u>apid <u>O</u>pensource <u>M</u>inimum Spanning Tre<u>E</u> Alg<u>O</u>rithm. ROMEO: i) uses up to three weights calculated from phase and magnitude information to provide improved unwrapping paths compared to BEST PATH, ii) provides computationally efficient bookkeeping of quality values and respective voxel edges, and iii) offers single-step unwrapping up to a fourth dimension (echo or time). We tested ROMEO’s performance against PRELUDE and BEST PATH in simulated topographies, challenging human head images acquired at 3 T and 7 T, and rat head images at 9.4 T. We provide compiled versions of ROMEO for Linux and Windows and source code in the Julia (17) programming language, all downloadable from: https://github.com/korbinian90/ROMEO. The complete datasets used in this study including the unwrapped phase images are also available at: https://dataverse.harvard.edu/dataverse/ROMEO.

## METHODS

### A. The ROMEO algorithm

It is important for the accuracy of path-based phase unwrapping methods that unwrapping proceeds along a path connecting reliable (albeit wrapped) voxels, as errors are introduced when noise is encountered. For this reason, it is crucial to make an astute choice of the order in which voxels are unwrapped, defined by the quality of the connection between them. ROMEO uses up to three weights which are multiplied together to yield a map of the “quality” of connection between neighbouring voxels for each of the 3 principal directions (x, y and z), here called a quality map. The unwrapping process is guided through the 3-dimensional phase data by this quality map, starting at a seed voxel with the highest quality value. For computational efficiency, the real-valued quality values are converted into integer cost values which are sorted into a bucket priority queue (18): a sequence of non-negative integers, each of which has a “priority” associated with it: the cost value ranking. This effectively creates a minimum spanning tree of the cost values of the edges during the unwrapping process according to the Prim-Jarník Algorithm (19). ROMEO was written in the open-source programming language Julia (17), which has a syntax of similar simplicity to that of Python and a speed similar to the C-based languages.

The ROMEO weights are defined in the range [0 ; 1] with ‘good’ weights (those indicating well-connected voxels) being close to 1, which allows easy combination of all or only some of the weights via multiplication. The three weights, calculated in each direction (x, y and z), are defined as follows:

1. Spatial phase coherence weight:

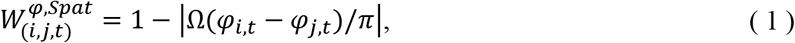

where Ω is a wrapping operator, *φ_i,t_* and *φ_j,t_* are measured phases at two adjacent spatial locations, *i* and *j*, and 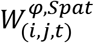 is the spatial weight of edge (*i,j*), all at the same time point *t*.
2. Temporal phase coherence weight:

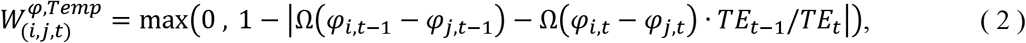

where *t* = 2 and phase values for the first and second echo (*TE_1_, TE_2_*) are chosen as the default for the calculation of the temporal coherence weight, 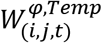. It is possible to change *t* if desired.
3. Magnitude coherence weight:

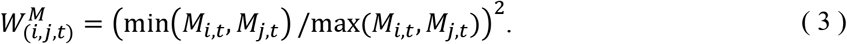

Weight 2 is only used for multi-echo or multi-time-point data and can be omitted. Weight 3 is used if magnitude data are available. An example of the three weights for the x-direction (left-right) for a 7 T 3D GRE dataset is shown in Figure 1.

**Fig. 1.**
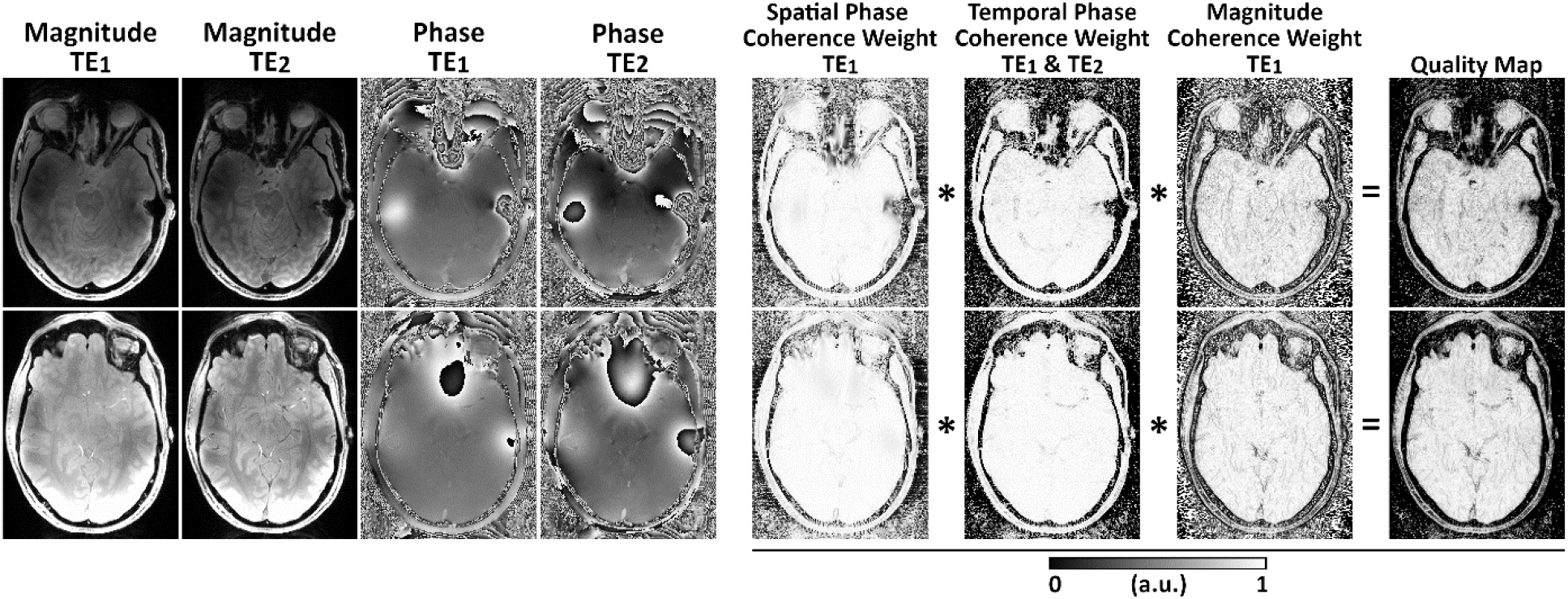
An example of maps of the three ROMEO weights for the x-direction (left-right): Magnitude and Phase images at the first two echo times, Spatial Phase Coherence, Temporal Phase Coherence, and Magnitude Coherence weights. Multiplication of these weights defines the final Quality Map for the x-direction. Two axial slices are shown from data acquired at 7 T with the 3D GRE sequence parameters shown in Table 1. These maps were calculated for the first echo using the magnitude image acquired at TE_1_ and phase images acquired at TE_1_ and TE_2_. The weights and quality values close to 1 correspond to good voxel connections; values close to 0 are weakly connected and are unwrapped last. Analogous weights are calculated for the y and z directions.

The product of the weights for each direction yields a quality map for each direction. For computational efficiency and reduced memory usage, the real valued quality map between 0 and 1 is transformed into integer cost values between 255 and 1, i.e. *cost* = max (*round*[255 · (1 − *quality*)), 1). An integer cost value of 1 corresponds to the best connection and a cost value of 255 to the worst. The special case of a cost value of 0 denotes no connection between voxels e.g. the border of a mask. In this case, the corresponding phase values are not included in the priority queue, which effectively stops the unwrapping process in that direction. The range of values from 0 to 255 was chosen as it can be efficiently stored as an 8 bit unsigned integer (2^8^ = 256) and represents the range of original real-valued quality values with sufficient accuracy to avoid changing the unwrapped result. Cost values derived from the quality map and corresponding voxel edge locations along the 3 different axes are passed into the bucket priority queue. The priority queue initially contains six cost values surrounding the seed voxel (in directions -x, x, -y, y, -z, z). The smallest value in the queue is identified together with the corresponding edge connecting the seed voxel (Voxel 1) with its neighbour (Voxel 2). If there is a phase jump ≥ π between these voxels, 2πn is subtracted from the phase of Voxel 2 according to: *θ*_2,*t*_ = *φ*_2,*t*_ − 2*πn* = *φ*_2*t*_ − 2*π* * 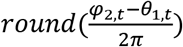, where *φ*_2,*t*_ is the wrapped phase measured in Voxel 2, *θ_1,t_* is the phase in Voxel 1 and *θ*_2,*t*_ is the unwrapped phase in Voxel 2, all at a given time point *t*. Voxel 2 is subsequently marked as having been visited. New values are added to the queue including the connections between Voxel 2 and all its neighbours not yet visited by the algorithm. When a new edge is drawn from the queue, a check is performed to see if the voxels connected by the edge have both been visited: if they have, this edge is removed from the queue. The search for the minimal cost value and the unwrapping process are repeated iteratively until all voxels have been visited.

By default, ROMEO calculates weights only for a single 3D volume in multi-echo or multi-time-point data - a template phase volume - and uses the unwrapped result from this template, *θ_i t_*, to unwrap the neighbouring volumes, *φ_i,t±i_*, assuming an approximately linear phase evolution in time:

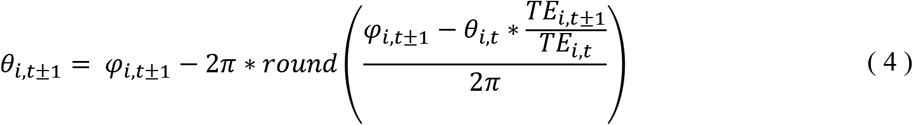

This accelerates the unwrapping process substantially by avoiding the recalculation of the weights for each single volume and improves the stability of the unwrapping results over the echoes or time points. By default, *t*=2 is specified as the template phase volume because *t*=1 tends to be more affected by flow effects (in multi-echo GRE) or is acquired prior to the longitudinal magnetisation reaching a steady state (in EPI time-series). The template phase volume can be changed if necessary. In specific cases, when large motion occurs between the time points or the assumption of linear phase evolution is not fulfilled, individual phase unwrapping can be applied with the calculation of weights and spatial unwrapping for each volume.

### B. Datasets

To provide a ground truth phase, θ, for a complicated pattern of wraps in φ, a complex topography was simulated as in Refs [8] and [10], with matrix size 256×256×256 and echo times TE = [4, 6, 10] ms. Example magnitude and phase images of this topography at TE = 10 ms are shown in Figure 2.

**Fig. 2.**
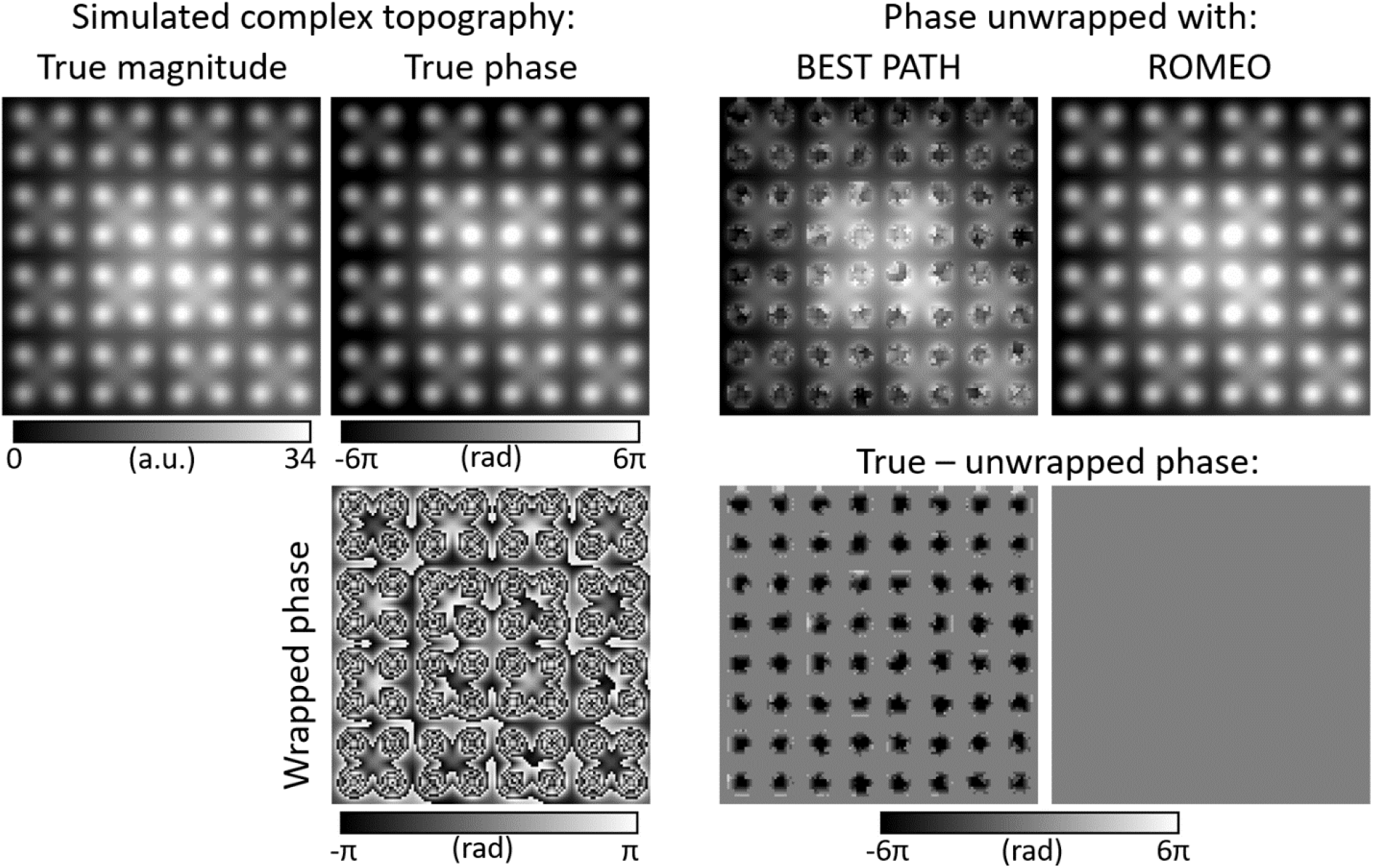
The BEST PATH and ROMEO phase unwrapping results for the simulated complex topography at TE = 10 ms. The difference between the true and the unwrapped phase shows no residual wraps with ROMEO unwrapping and many errors with BEST PATH unwrapping. PRELUDE failed to deliver results within 38 days.

Measured phase maps (with no ground truth θ) were also examined: in vivo human head MRI acquisitions at 3 T and 7 T (Siemens MAGNETOM, Siemens Healthineers, Erlangen, Germany) and ex vivo rat head images acquired at 9.4 T (Bruker BioSpec 94/20 USR, Ettlingen, Germany) with sequence parameters listed in Table 1.

**Table 1.**
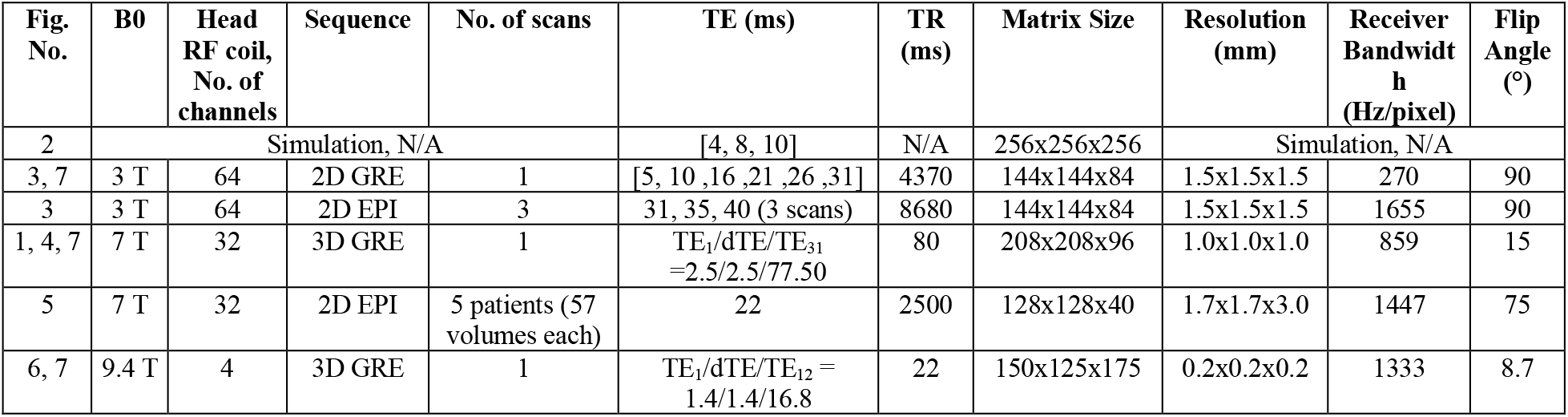
Datasets used for the assessment of ROMEO unwrapping accuracy and computational speed.

Human measurements were approved by the Ethics Committee of the Medical University of Vienna and all participants provided written informed consent. All human data sets were acquired from healthy volunteers except for five 7 T EPI time series (57 volumes) from a prior study (20), which were collected from four patients with brain tumours and one patient with a developmental venous anomaly.

### C. Data analysis

All the data were acquired with multi-channel coil arrays. Separate channels were combined using ASPIRE (21) for multi-echo GRE data and using the coil combination described in Ref. (22) for the single-echo EPI data. Combined phase images were unwrapped using compiled PRELUDE from the FSL toolbox (version 5.0.11, https://fsl.fmrib.ox.ac.uk/fsl/fslwiki/FSL) written in C++ (8), BEST PATH (9) programmed in C, and ROMEO written in Julia programming language. Their performance was compared with respect to unwrapping accuracy and computational speed. All the calculations were performed on a PC with an Intel Xeon W-2125 4.0GHz processor, 64GB RAM and an Ubuntu Linux 16.04 operating system.

All ROMEO results were obtained using magnitude and phase images as well as template unwrapping as described in the Methods section A. Unmasked measured data were analysed for BEST PATH and ROMEO. Images were additionally masked to obtain PRELUDE results in a feasible time. Masking was performed with the FSL Brain Extraction Tool (BET) (23) for in vivo data other than the 3 T EPI, where SPM Segment Toolbox (SPM12, https://www.fil.ion.ucl.ac.uk/spm/) was used, as BET produced masks which did not match the image well. Ex vivo rat head images were masked using magnitude image thresholding.

A qualitative comparison of the results was performed using MRIcro (https://www.mccauslandcenter.sc.edu/crnl/mricro) and FSLeyes (https://fsl.fmrib.ox.ac.uk/fsl/fslwiki/FSLeyes) and selected slices with substantial differences between the three phase unwrapping methods are presented in the Results section A.

For the simulated data, quantitative comparison was performed by calculating the percentage of unwrapped values which were different from the ground truth. Although for in vivo measurement there is inherently no ground truth available, a reliable estimate of the true phase, or Temporal Reference image, can be obtained for multi-echo data if the first echo time is short and the echo time difference between consecutive echoes is small, ensuring small and approximatively linear phase evolution between the echoes. This was found to be the case for the multi-echo GRE data at 3 T, 7 T and 9.4 T. For the first echo time, the Temporal Reference image was merged from the results of the three methods analysed here: voxels were only included if their unwrapped phase values were the same for all three unwrapping methods. This meant that for the 3 T, 7 T and 9.4 T GRE data respectively, 99%, 98% and 89% of the voxels within the brain mask were included in the Temporal Reference image. The Temporal Reference for subsequent echoes was calculated by assuming approximately linear phase evolution over time using Equation 4. Phase unwrapping errors were calculated as the difference between the unwrapped phase obtained using a given method at a given TE and the Temporal Reference at the same TE. Histograms showing the number of voxels with 2πn errors for all three methods analysed were plotted (see Figure 7). The percentage of voxels with unwrapped phase values different than the Temporal Reference within the mask is listed in Table 2.

**Table 2.** The percentage of erroneous voxels within the mask for all three phase unwrapping methods and second, middle and last echo of the GRE acquisitions at 3 T, 7 T and 9.4 T. Voxels with 2πn phase differences (where n is an integer) from the Temporal Reference were counted as erroneous (see Method section C).

|  |  | No. of erroneous voxels within the mask [%] |  |  |
| --- | --- | --- | --- | --- |
|  | TE [ms] | PRELUDE | BEST PATH | ROMEO |
| **3 T GRE** | 10.0 | 0.5301 | 0.6065 | 0.4168 |
|  | 21.0 | 1.7381 | 1.9225 | 0.4172 |
|  | 31.0 | 2.9734 | 3.3129 | 0.4174 |
| **7 T GRE** | 5.0 | 0.2700 | 0.2906 | 0.1155 |
|  | 40.0 | 5.7006 | 5.3991 | 0.1160 |
|  | 77.5 | 22.8113 | 23.3927 | 0.1217 |
| **9.4 T GRE** | 2.8 | 1.4462 | 1.7392 | 0.8535 |
|  | 9.8 | 14.9638 | 13.9962 | 0.8570 |
|  | 16.8 | 26.5759 | 24.8455 | 0.8633 |

For EPI at 3 T and 7 T, calculation of a Temporal Reference as described above was not possible due to the inherently long minimum echo times of the EPI acquisitions. These images were assessed visually, using MRIcro and FSLeyes. For the single-echo 7 T EPI time-series data, temporal mean and standard deviation (SD) images were calculated throughout the brain mask to investigate regions where unwrapping errors were different at various time points (see Figure 5). The SD of the estimated field map 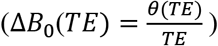 was calculated for the 9.4 T GRE dataset with 12 echoes (see Figure 6).

## RESULTS

### A. Comparison of unwrapping accuracy between PRELUDE, BEST PATH and ROMEO

Figure 2 shows BEST PATH and ROMEO unwrapping results for the simulated topography data at TE = 10 ms. PRELUDE failed to complete unwrapping within 38 days (912 hours). BEST PATH delivered accurate results only for TE = 4 ms. At TE = 8 ms 216 voxels, (0.001% of all voxels) and at TE = 10 ms 1262372 voxels (7.524% of all voxels) in the BEST PATH results had incorrect phase values. The difference image between the true and the unwrapped phase highlights errors in the BEST PATH results at TE = 10 ms. ROMEO provided an accurate outcome, leaving no residual wraps at all echoes.

Unwrapping results for the 3 T GRE and EPI dataset with TE = 31 ms are presented in Figure 3. Two slices with visible differences between PRELUDE, BEST PATH and ROMEO are shown. Unwrapping errors occurred in all methods close to the sinuses (red arrows), where BEST PATH shows the largest errors. In both the GRE and EPI phase, an open-ended fringe line is clearly visible in the wrapped phase close to the left ear canal where the magnitude signal approaches the noise level (blue arrows). In ROMEO, the extent of the unwrapping error in this region was limited to a few voxels. In PRELUDE and BEST PATH, the size of the region affected increased with echo time (not shown). A residual wrap also occurred in the vein of Galen in the BEST PATH result (see Figure 3 GRE, yellow arrows). Unwrapping differences between the three methods were also observed in a small number of voxels in other vessels (see Figure 3 EPI, yellow arrows). Regions affected by residual wraps were the smallest in ROMEO.

**Fig. 3.**
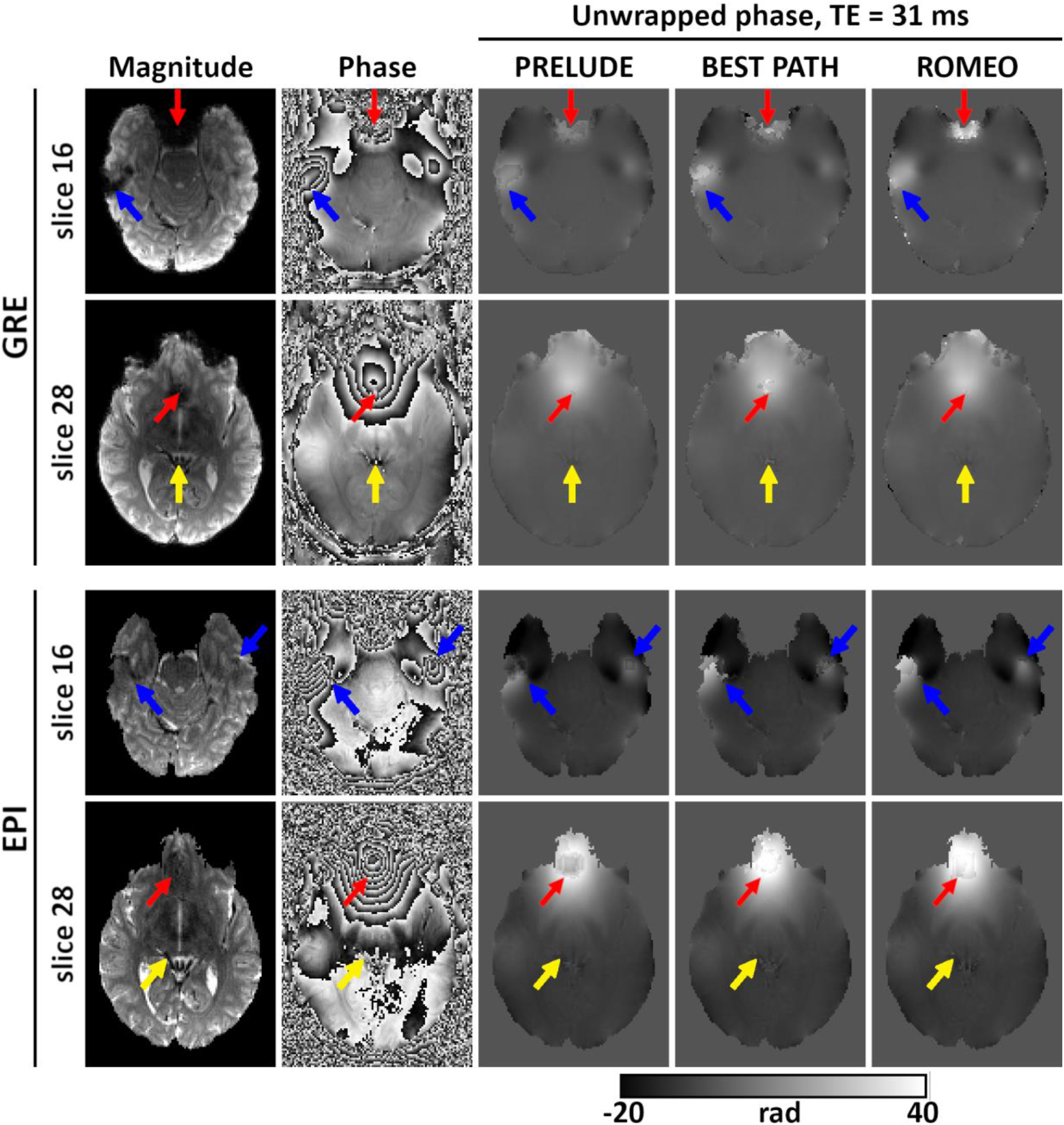
Unwrapping results for the 3 T GRE and EPI scans at TE = 31 ms. Magnitude and phase images (wrapped and unwrapped) are shown for two slices. Red arrows point to the regions close to the sinuses, blue arrows point to open-ended fringe lines close to ear canals and yellow arrows pointto vessels affected by low signal and unwrapping errors. Overall, ROMEO results had the smallest number of voxels with residual phase wraps.

Examples of phase unwrapping performance for PRELUDE, BEST PATH and ROMEO at 7 T are presented in Figure 4 for multi-echo GRE and in Figure 5 for a single-echo EPI time series. A central axial slice from the GRE data is shown at four selected echo times in Figure 4 starting with a relatively short echo time (TE_6_ = 15 ms) and ending with a very long one (TE_30_ = 75 ms), where the signal in a large part of the image has decayed into noise. At TE_6_ = 15 ms, small differences were observed at the brain boundaries and in the sagittal sinus (red arrows) between the Temporal Reference and the PRELUDE or BEST PATH results. At TE_15_ = 30 ms slightly larger regions with unwrapping errors were observed in the PRELUDE and BEST PATH results, for example close to the auditory canals (red arrows). At later echoes, such as TE_24_ = 60 ms and TE_30_ = 75 ms, large patches of tissue were affected by phase unwrapping errors in both the PRELUDE and BEST PATH results with larger regions of errors observed at longer TEs. No difference between the Temporal Reference and the ROMEO unwrapped phase is visible in this slice at TE_6_ and TE_15_, but differences in a few voxels are observed at TE_24_ and TE_30_ (blue arrows). Regions with a very low SNR, approaching the noise floor in the magnitude image, are noisy in both the Temporal Reference and the ROMEO unwrapped phase, otherwise both show a coherent phase topography. Small differences between the two results are more apparent in the quantitative comparison of the methods in Figure 7 and Table 2.

**Fig. 4.**
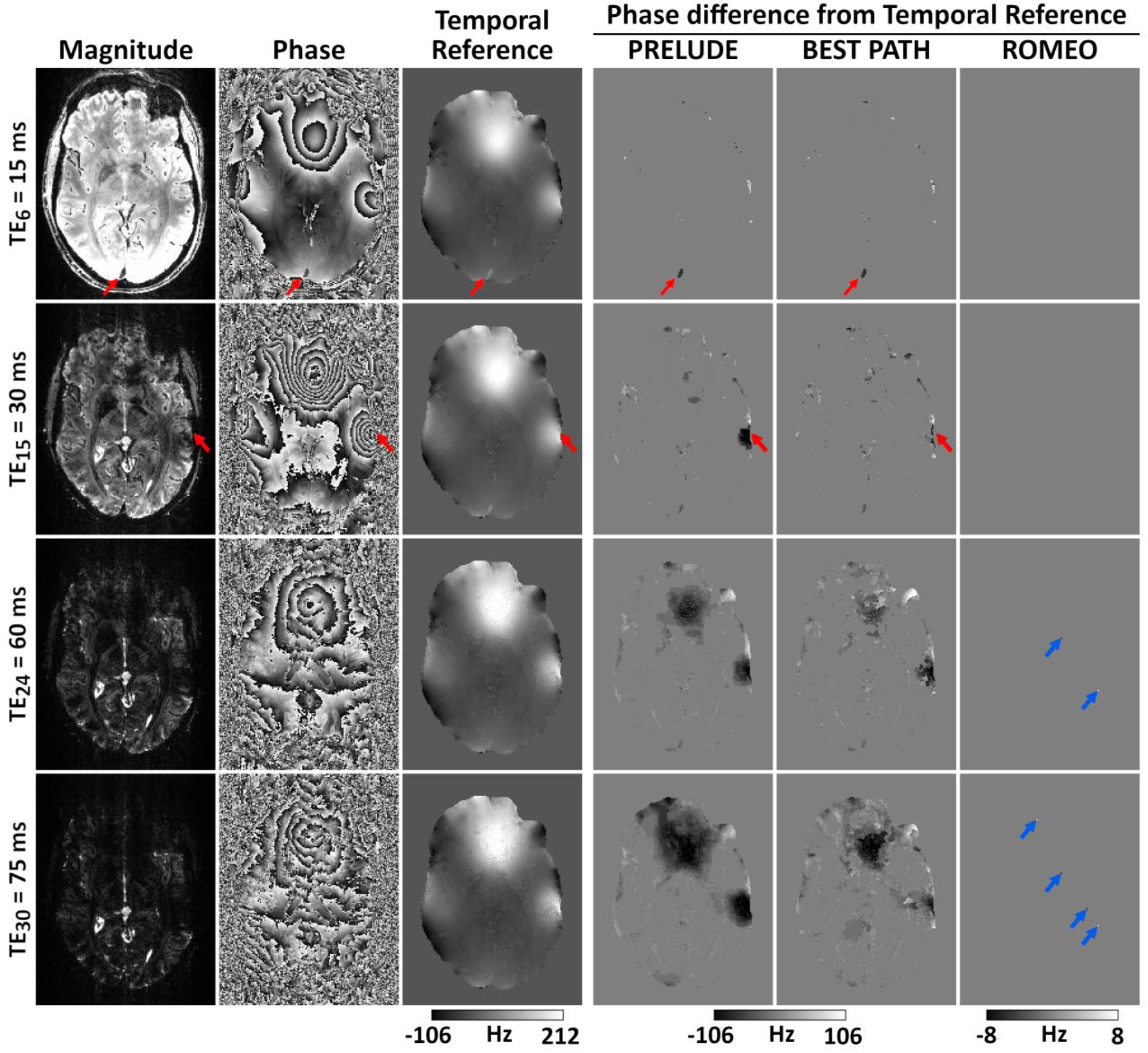
Unwrapping results for the 7 T GRE data at four selected echoes (of 31). At shorter echoes (TE_6_ = 15 ms and TE_15_ = 30 ms) a few unwrapping differences between the PRELUDE or BEST PATH and the Temporal Reference phase occur at the brain edges, sagittal sinus or ear canals (see red arrows). At longer echoes (TE_24_ = 60 ms and TE_30_ = 75 ms) large patches with unwrapping errors are visible in PRELUDE and BEST PATH results. There is no difference between the Temporal Reference phase and the ROMEO result at TE_6_ and TE_15_, and only a few voxels (marked by blue arrows) differ at longer echoes.

**Fig. 5.**
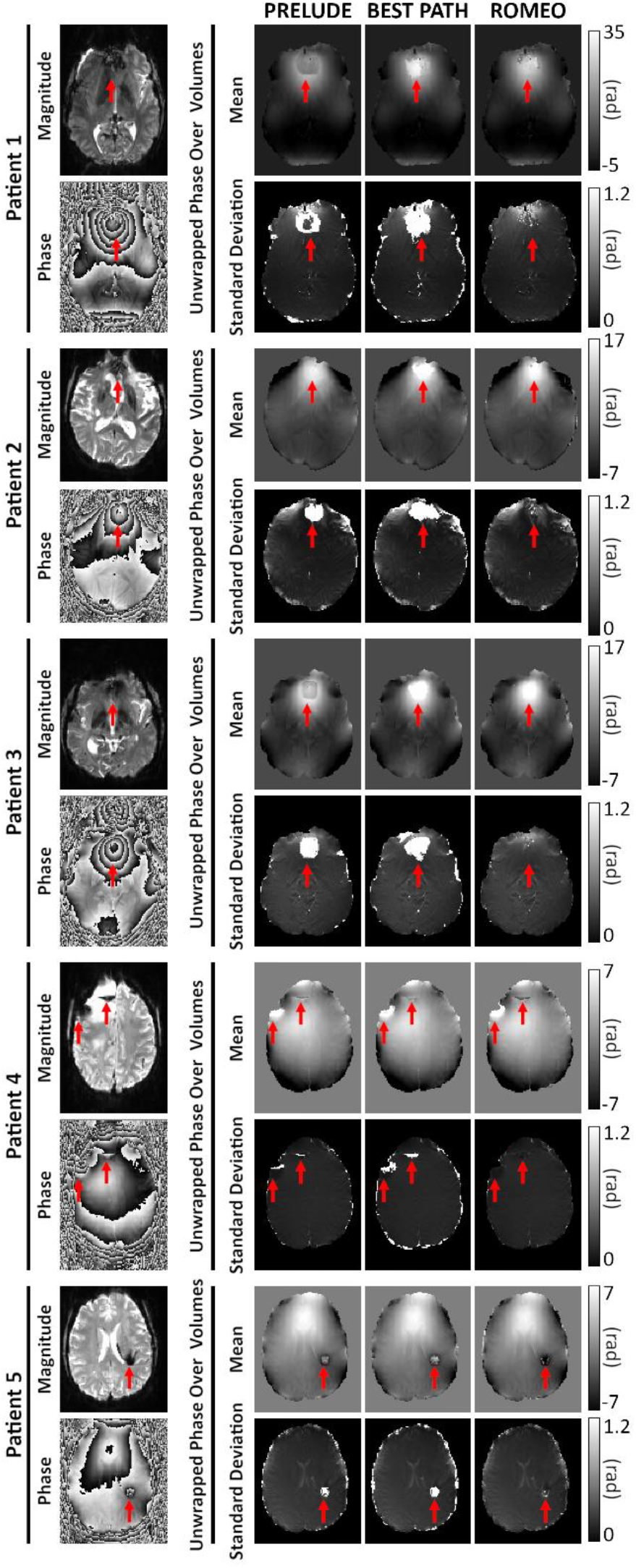
Unwrapping results for a 7 T EPI time series with 57 volumes acquired in four patients with brain tumours and one patient (Patient 5) with a developmental venous anomaly. The temporal mean and standard deviation of the unwrapped phase are shown for all patients. Standard deviation maps highlight residual phase wraps which change for different time points. Red arrows highlight the largest errors. ROMEO outperformed PRELUDE and BEST PATH, yielding both fewer residually wrapped voxels and less temporal variation in the unwrapped phase over time points.

PRELUDE and BEST PATH results for the 7 T EPI time series (57 volumes) were affected by global 2πn phase jumps between consecutive time points, numbering 0, 19, 0, 14 and 32 jumps respectively for patients 1 to 5 for PRELUDE and 17, 10, 18, 0 and 26 for BEST PATH. No phase jumps between time points were present in the ROMEO results. Therefore, global phase jumps in the PRELUDE and BEST PATH results were removed prior to the calculation of the temporal mean and standard deviation of the unwrapped phase, which are shown in Figure 5. In patients 1, 2 and 3, extensive unwrapping errors occurred using PRELUDE and BEST PATH close to the sinuses, marked by the arrows. These errors changed in size at different time points, which contributed to the high values of the phase standard deviation in these regions. ROMEO unwrapping errors were substantially smaller and stable over time points, which is reflected in low standard deviation values. In patients 4 and 5, the unwrapping errors occurred close to pathologies and were, again, less extensive for ROMEO than for PRELUDE or BEST PATH.

Figure 6 shows high resolution images of an ex vivo rat brain, acquired at 9.4 T using a multi-echo GRE sequence. ROMEO gave the most accurate phase unwrapping results, agreeing well with the Temporal Reference, and with very good stability over the echoes, as highlighted by the Δ*B*_0_ SD. Regions in PRELUDE and BEST PATH results marked by red arrows were affected by residual phase errors, which changed for different echoes (see corresponding Δ*B*_0_ SD maps). Blue arrows point to the superior sagittal sinus, which had a phase offset with respect to the surrounding tissue in both the Temporal Reference and ROMEO results, which was consistent for all echoes, as represented by the low Δ*B*_0_ SD values. The phase images unwrapped by PRELUDE and BEST PATH show similar offsets in the sagittal sinus but only at some of the shorter echo times (5 echoes in PRELUDE and 7 in BEST PATH), which is reflected by high Δ*B*_0_ SD values in this large vein.

**Fig. 6.**
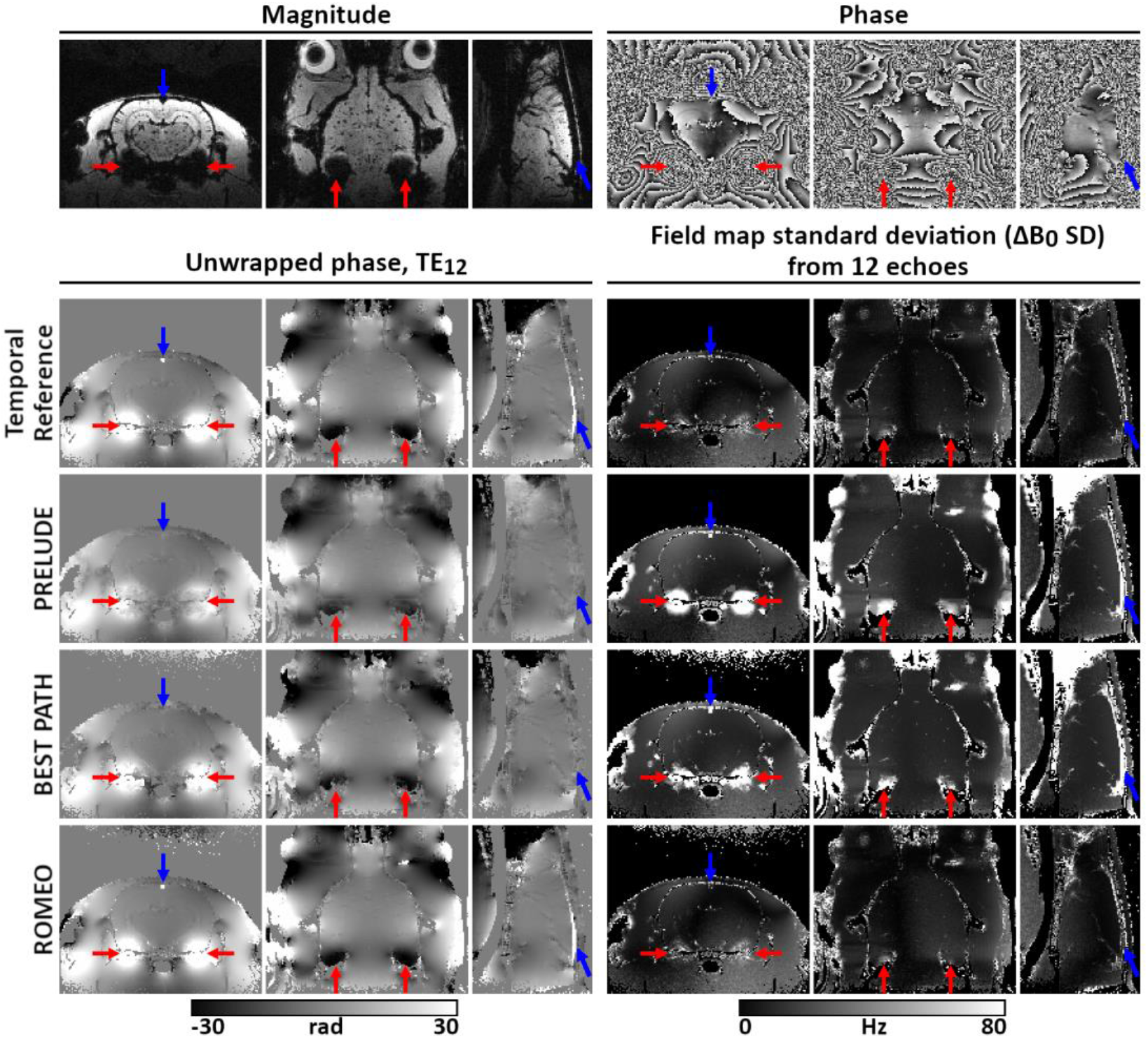
Unwrapping results for images of a rat brain acquired at 9.4 T using a multi-echo GRE sequence (TE_1_ / dTE / TE_12_ = 1.4 / 1.4 / 16.8 ms). The magnitude and unwrapped phase from the last echo (TE_12_) and the field map standard deviation (Δ*B*_0_ SD) over all 12 echoes are shown in all three perpendicular planes. Differences between the methods are highlighted by red and blue arrows. ROMEO unwrapping results are the most accurate, the most similar to the Temporal Reference and the most stable over echoes.

Histograms showing the number of voxels with 2πn phase errors (n is an integer) in the unwrapped GRE results at 3 T, 7 T and 9.4 T, for the middle echo, the last echo, and over all echoes are presented in Figure 7. PRELUDE and BEST PATH show similar error spectra, which increase in amplitude and become broader at longer echo times. The number of erroneous voxels in ROMEO is substantially smaller than for the other unwrapping methods. This result is also highlighted in Figure 7, where the percentage of erroneous voxels within the mask is listed for all methods and three selected echoes. For all methods, the number of erroneous voxels increased with the TE. For ROMEO results this number stayed below 1% at all field strengths and all echoes. PRELUDE and BEST PATH errors affected over 20% of the voxels at the longest echoes at 7 T and 9.4 T.

**Fig. 7.**
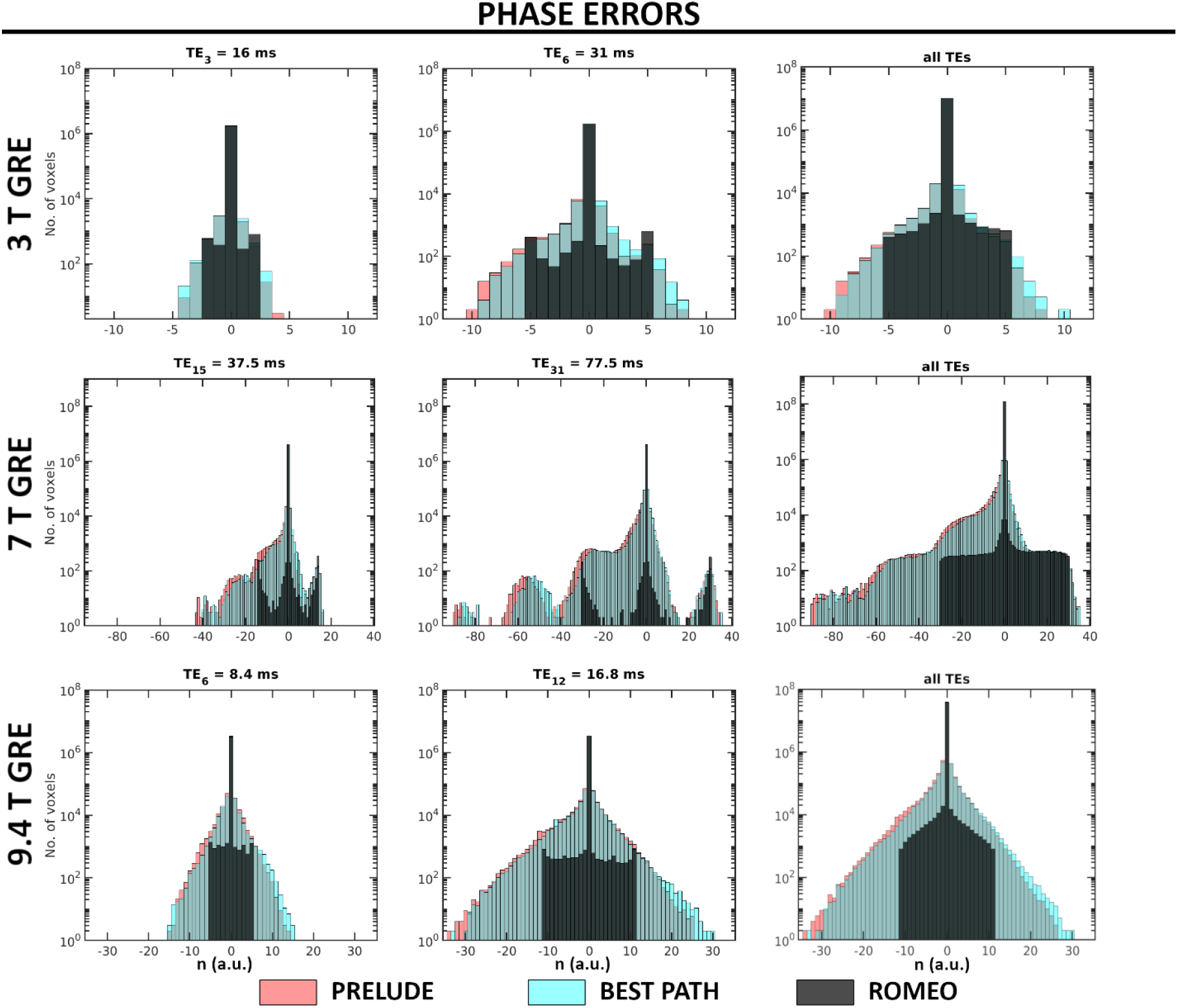
Histograms of phase errors in PRELUDE, BEST PATH and ROMEO GRE results at all field strengths for the central and last echo times, and summed over all echoes (excluding TE_1_). Voxels with 2πn (where n is an integer) phase differences from the Temporal Reference phase were counted as erroneous (see Method section C). The number of voxels is shown on a logarithmic scale.

All wrapped phase images described above were masked to obtain PRELUDE results in feasible times. For BEST PATH and ROMEO no mask was required, therefore unwrapping with no mask was also assessed. This yielded identical results, within masks, to the analysis above. There was, however, a difference in computation time between executions with or without a mask, as described below.

### B. Comparison of computational speed between PRELUDE, BEST PATH and ROMEO

The computation times of ROMEO in comparison to PRELUDE and BEST PATH are summarized in Table 3. PRELUDE unwrapping took from several minutes (14 min 36 sec for 3T GRE) to several hours (128 hours 8 min 59 sec for 7 T GRE) and the unwrapping process failed to finish within 38 days for the simulated dataset. BEST PATH took less than a minute for all masked data, and ROMEO less than 30 seconds. ROMEO was generally faster than BEST PATH with an exception for 3 T datasets, where BEST PATH was faster by at most 4 seconds (see Table 3 for more details).

**Table 3.** Computation times for PRELUDE, BEST PATH and ROMEO. The fastest times for each dataset are in bold.

|  |  | Computation time (hh:mm:ss) |  |  |
| --- | --- | --- | --- | --- |
| Dataset |  | PRELUDE | BEST<br>PATH | ROMEO |
| Simulation | Not masked | Not<br>finished<br>in 912 h | 00:00:38 | 00:00:20 |
| 3 T GRE | Masked<br>Not masked | 00:14:36 | 00:00:04<br>00:00:07 | 00:00:06<br>00:00:07 |
| 3 T EPI | Masked<br>Not masked | 00:32:25 | 00:00:02<br>00:00:04 | 00:00:06<br>00:00:07 |
| 7 T GRE | Masked<br>Not masked | 128:08:59 | 00:00:48<br>00:00:91 | 00:00:09<br>00:00:11 |
| 7 T EPI |  |  |  |  |
| Patient 1 | Masked<br>Not masked | 01:03:29 | 00:00:11<br>00:00:26 | 00:00:06<br>00:00:06 |
| Patient 2 | Masked<br>Not masked | 00:30:37 | 00:00:11<br>00:00:25 | 00:00:06<br>00:00:06 |
| Patient 3 | Masked<br>Not masked | 00:29:03 | 00:00:11<br>00:00:25 | 00:00:06<br>00:00:06 |
| Patient 4 | Masked<br>Not masked | 00:38:43 | 00:00:09<br>00:00:26 | 00:00:06<br>00:00:06 |
| Patient 5 | Masked<br>Not masked | 00:27:22 | 00:00:11<br>00:00:26 | 00:00:06<br>00:00:07 |
| 9.4 T GRE | Masked<br>Not masked | 56:03:07 | 00:00:17<br>00:00:27 | 00:00:08<br>00:00:09 |

Using BEST PATH and ROMEO, datasets that were not masked took longer to unwrap than masked images. This difference was less prominent for ROMEO, with the same unwrapping times for masked and unmasked 7 T EPI datasets (6 seconds). All three methods were memory-efficient with the maximum RAM usage below 5 GB for datasets (magnitude and phase images) size below 800 MB.

## DISCUSSION

We have presented a new, rapid and robust phase unwrapping technique – ROMEO – that is more reliable and faster than the two exact phase unwrapping algorithms most commonly used in MRI; PRELUDE and BEST PATH. Since the MRI signal is complex, magnitude information is available for every MRI scan, even if a study focuses exclusively on phase imaging. ROMEO includes information about the spatial coherence of the magnitude signal in the unwrapping operation. Combining this information with information on the phase’s spatial and temporal coherence creates a refined quality map that guides the unwrapping process through three-dimensional data, starting with the most reliable voxels. This improves the unwrapping accuracy over BEST PATH, which uses a quality map based on only a single weight calculated from the second difference of the phase between the six nearest neighbours and the twenty diagonal neighbours of a given voxel. Moreover, ROMEO uses ‘template-based’ unwrapping with respect to a selected template volume when the data have a fourth dimension (e.g. echoes or time points). This allows ROMEO to avoid introducing 2πn phase jumps between these echoes or time points and speeds up the unwrapping operation, as the weights and quality map are calculated only for one 3D template volume. Template unwrapping works accurately if the phase varies approximately linearly with the echo time and no residual phase offsets are present (i.e. θ ≈ 0 at TE = 0). If this is not the case (e.g. due to large motion) ROMEO offers the option of unwrapping each volume individually.

The phase unwrapping problem exists since it is not possible to measure the ground truth phase, which means that creating a reference image and performing a quantitative comparison of different phase-unwrapping algorithms in vivo is very challenging. The Temporal Reference was calculated with the assumption that the phase evolves approximately linearly over time, which is, for the purpose of assessing wraps, a reasonable approximation of the ground truth. It is only possible to calculate such a Temporal Reference from multi-echo acquisitions with a short initial echo time and small echo spacings. The first echo-time phase must be either free of wraps or unwrapped without errors by all the methods under evaluation. In addition to a thorough qualitative comparison presented in Figures 2–6, we have also provided a quantitative analysis of unwrapping errors for all three phase unwrapping methods considered here: see Figure 7 and Table 2. We have calculated the percentage of voxels in each method with values different from the ground truth in simulation or from the Temporal Reference in measured data. ROMEO uses template unwrapping, a type of temporal unwrapping, which yielded results that agreed well with the Temporal Reference.

Path-following methods, BEST PATH and ROMEO, were much faster than PRELUDE, a region-growing method. The improved speed of ROMEO with respect to BEST PATH arises from template unwrapping as well as from efficient handling of values in the queue of voxels to be considered. BEST PATH deploys the Kruskal algorithm to calculate the Minimum Spanning Tree (MST) (24), using a heap as the priority queue, which has a runtime that depends on the number of voxel insertions into the queue, m, according to O(m log(m)). ROMEO uses the Prim-Jarník Algorithm (19) with integer representation in a bucket priority queue, the runtime of which scales with O(m). Some of the speed differences may also come from the fact that the two methods were implemented in different programming languages (BEST PATH in C, ROMEO in Julia).

ROMEO took only a few seconds even for very challenging examples such as 7 T GRE images (9 seconds for masked images), where PRELUDE delivered results after about 128 hours and BEST PATH took 48 seconds. Although ROMEO was usually several seconds faster than BEST PATH (except in 3T datasets), this is less relevant than the unwrapping accuracy. ROMEO showed fewer residual phase wraps than BEST PATH and PRELUDE in all the cases analysed. Additionally, ROMEO demonstrated superior phase unwrapping stability over time points or echoes, which was highlighted by phase or field map standard deviation results. PRELUDE and BEST PATH often showed a different distribution of residual wraps in problematic areas (e.g. close to open-ended fringe lines) at different time points, rendering large areas of the phase images unusable, and requiring post-hoc global 2πn jump correction between adjacent volumes.

The unwrapping accuracy was independent of masking for BEST PATH and ROMEO, which highlights the redundancy of masking for these path-following methods and allows time to be saved that is normally spent on the often-fraught problem of mask generation. PRELUDE only generated results where a mask was provided but even then, calculation times were excessively long. As shown using simulated data, not masking PRELUDE inputs led to the algorithm failing to yield results even after many days. The quality maps calculated in ROMEO can be combined and thresholded to generate an object mask. This could be useful in applications requiring a mask such QSM, particularly for inhomogeneous images and non-brain regions or phantoms, where commonly used methods such as BET (23) do not perform well. ROMEO is extremely flexible as it has the option to output individual weights and the quality map (both combined over x, y and z), as well as a mask.

There is substantial interest in using EPI sequences for phase imaging and QSM (25–29). Phase images from EPI acquisitions were generally more challenging to unwrap than those from GRE scans because EPI has a lower SNR than GRE and is affected by other effects such as distortions in the phase encoding direction or stronger eddy currents. Of the three methods tested, ROMEO proved to be the most accurate and robust unwrapping algorithm for single-echo EPI acquisitions.

Three weights were included in the ROMEO implementation discussed here. We offer the source code in the Julia programming language which allows users to experiment with alternative weights for atypical MRI acquisitions or phase datasets acquired using other modalities such as optical or satellite radar interferometry.

We expect ROMEO to find applications in MRI phase imaging and QSM, especially in challenging cases such as: at high fields, at long echo times, in highly accelerated datasets with low SNR, and close to air spaces or implants. ROMEO’s speed makes it feasible to use spatial phase unwrapping in real time on the MRI reconstruction computer, which could benefit large studies with hundreds or thousands of phase volumes, including functional MRI studies where phase information can be used to correct distortions (22,30) and provide quantitative information about changes in blood susceptibility in functional QSM (27–29).

## CONCLUSION

We have developed a new path-following phase unwrapping algorithm called ROMEO, which is more accurate and faster than PRELUDE and BEST PATH, yielding results within a few seconds even for highly wrapped data with large matrix sizes. ROMEO does not require explicit masking and allows single-step unwrapping of multi-echo or multi-time-point data with excellent stability over volumes. Therefore, it is suitable for MRI studies where phase unwrapping is challenging e.g. at high fields, with implants or in large datasets including functional MRI studies.

## ACKNOWLEDGEMENTS

This study was funded by the Austrian Science Fund project FWF31452. B. Dymerska and S. Robinson were supported by Marie Skłodowska-Curie Actions MRI COMIQSUM 798119 and MS-fMRI-QSM 794298 respectively and K. Shmueli by ERC Consolidator Grant DiSCo MRI SFN 770939. Additional support from the Austrian Federal Ministry for Digital and Economic Affairs and the National Foundation for Research, Technology and Development is gratefully acknowledged.

## DATA AVAILABILITY STATEMENT

The code that supports the findings of this study is openly available in GitHub at https://github.com/korbinian90/ROMEO and data in Harvard Dataverse at https://dataverse.harvard.edu/dataverse/ROMEO.

